# Streamlined single-molecule RNA-FISH of core clock mRNAs in clock neurons in whole mount *Drosophila* brains

**DOI:** 10.1101/2022.09.05.506677

**Authors:** Ye Yuan, Marc-Antonio Padilla, Dunham Clark, Swathi Yadlapalli

## Abstract

Circadian clocks are ∼24-hour timekeepers that control rhythms in almost all aspects of our behavior and physiology. While it is well known that subcellular localization of core clock proteins plays a critical role in circadian regulation, very little is known about the spatiotemporal organization of core clock mRNAs and its role in generating ∼24-hr circadian rhythms. Here we describe a streamlined single molecule Fluorescence *In Situ* Hybridization (smFISH) protocol and a fully automated analysis pipeline to precisely quantify the number and subcellular location of mRNAs of *Clock*, a core circadian transcription factor, in individual clock neurons in whole mount *Drosophila* adult brains. Specifically, we used ∼48 fluorescent oligonucleotide probes that can bind to an individual *Clock* mRNA molecule, which can then be detected as a diffraction-limited spot. Further, we developed a machine learning-based approach for 3-D cell segmentation, based on a pretrained encoder-decoder convolutional neural network, to automatically identify the cytoplasm and nuclei of clock neurons. We combined our segmentation model with a spot counting algorithm to detect *Clock* mRNA spots in individual clock neurons. Our results demonstrate that the number of *Clock* mRNA molecules cycle in large ventral lateral clock neurons (lLNvs) with peak levels at ZT4 (4 hours after lights are turned on) with ~80 molecules/neuron and trough levels at ZT16 with ∼30 molecules/neuron. Our streamlined smFISH protocol and deep learning-based analysis pipeline can be employed to quantify the number and subcellular location of any mRNA in individual clock neurons in *Drosophila* brains. Further, this method can open mechanistic and functional studies into how spatiotemporal localization of clock mRNAs affect circadian rhythms.

## Introduction

Almost all organisms have evolved internal clocks to tell time. Circadian clocks are cell-autonomous timekeepers that can be entrained by light^1^ or temperature^2-4^ cycles. These clocks generate ∼24-hour oscillations in the expression of many genes and control rhythms in sleep-wake cycles, metabolism, immunity^5-8^. While past studies highlight the critical role of subcellular localization of core clock proteins in controlling circadian rhythms^9-14^, what remains poorly understood is how clock mRNAs are organized over the circadian cycle and the role of such organization in controlling circadian rhythms. A major drawback of techniques such as qPCR and single-cell RNA-sequencing currently used to study circadian gene expression is lack of subcellular spatial information^15,16^. Single-molecule RNA Fluorescence *In Situ* Hybridization (smFISH) has recently emerged as a powerful technique which enables quantitative measurement of the number as well as the subcellular location of mRNA molecules in individual cells^17,18^. In one recent study, smFISH was adapted to visualize core clock transcripts in *Neurospora crassa* and it was reported that core clock mRNAs are clustered in the perinuclear cytoplasmic region in a time-of-day dependent manner^19^.

*Drosophila melanogaster* is a powerful model system for understanding spatiotemporal regulation of circadian rhythms, as it has a highly conserved clock along with powerful genetic and molecular tools^8,20^. Further, the *Drosophila* clock network is relatively simple consisting of ∼150 clock neurons^20,21^ (**Figure 1A**). These clock neurons express the key clock proteins– CLOCK (CLK), CYCLE (CYC), PERIOD (PER) and TIMELESS (TIM), that form the core of the molecular clock^8^ (**Figure 1B**). CLK and CYC are the positive transcription factors and PER and TIM act as the core clock repressors. During the activation phase, CLK and CYC drive transcription of hundreds of target genes, including *per* and *tim*, by binding to the E-boxes in their promoter regions. PER and TIM, after a time delay, enter the nucleus and inhibit CLK/CYC activity thereby silencing their own expression as well as the expression of other clock-regulated genes. PER and TIM are then degraded leading to the end of the repression phase and the start of the activation phase of a new cycle^8^. In addition to this core feedback loop, there exists a second interlocked feedback loop that controls rhythmic expression of *Clk* mRNA^22,23^ (**Figure 1B**).

**Figure 1.**
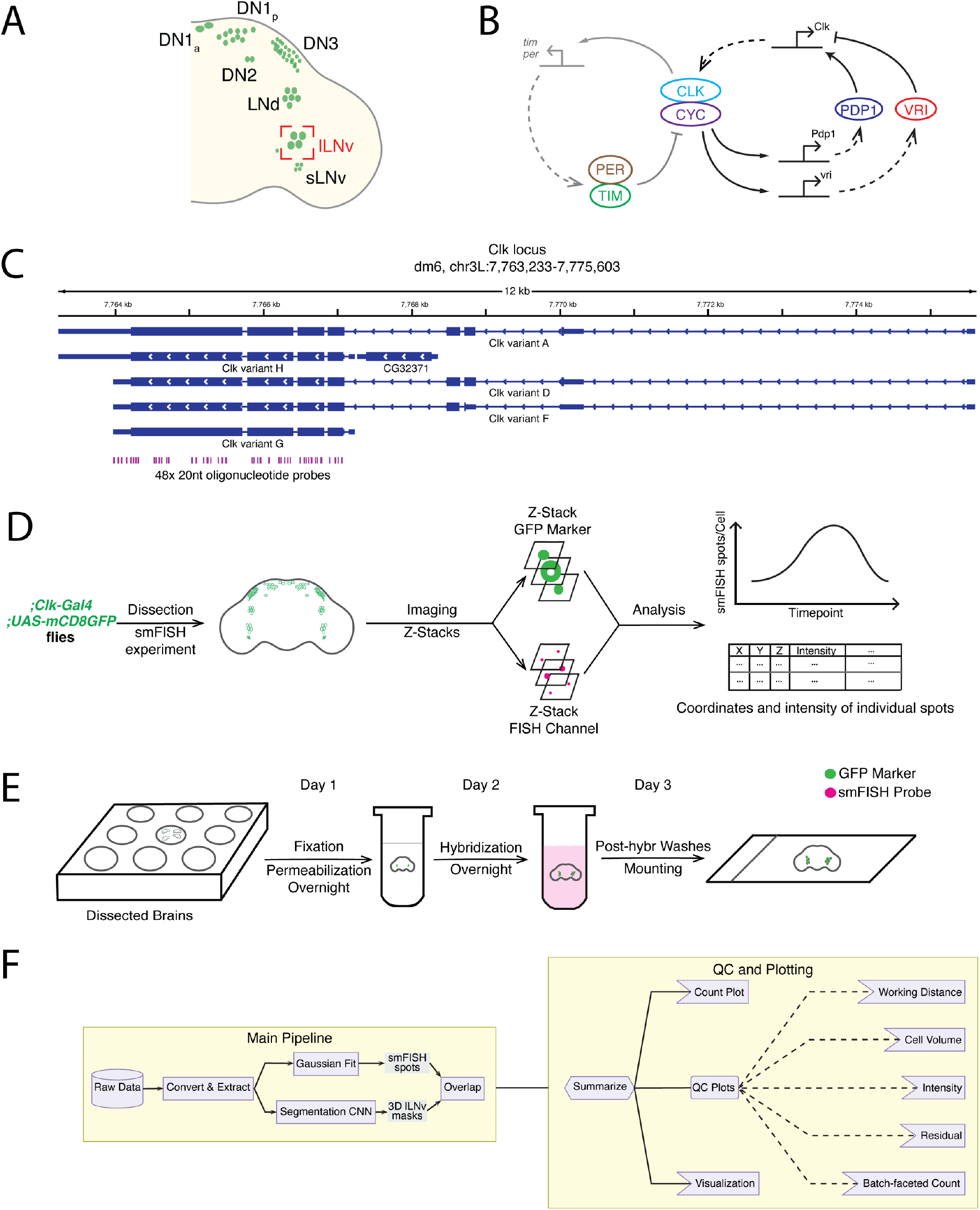
Overview of the smFISH method and the data analysis pipeline. **A)** Schematic of different clock neuron groups in *Drosophila melanogaster* brains. Large ventrolateral neurons (lLNvs) are circled in red. **B)** Schematic of the molecular clock in *Drosophila*. The *Drosophila* circadian clock is composed of two interlocked negative feedback loops. **C)** Genomic locations of oligonucleotide probes in the *Clock* locus. Probes are designed to hybridize with the shared exon sequences to visualize mRNAs of all annotated *Clock* transcript variants. **D)** Overview of the full workflow including smFISH experiment and data analysis pipeline. **E)** Schematic of the smFISH procedure. **F)** Schematic of the data analysis pipeline. To quantify *Clock* smFISH spots in lLNvs, two tasks are performed in parallel: detection of *Clock* smFISH spots and segmentation of lLNvs.

CLK/CYC proteins bind to the E-boxes of the transcription factors, Vrille (VRI) and PAR domain protein 1ε (PDP1ε), creating rhythms in their mRNA with peak levels during early night and trough levels during early day. Interestingly, VRI and PDP1ε protein levels peak at different times during the night, with VRI peaking during the early evening and PDP1ε later in the night. VRI and PDP1ε proteins bind one after the other to the VRI/PDP1ε-binding boxes in the *Clk* enhancer to repress or activate *Clk* transcription, respectively, thus resulting in rhythmic *Clk* mRNA expression.

Here, we describe a streamlined single molecule FISH (smFISH) protocol and a fully automated analysis pipeline to quantify the number and localization of mRNAs of *Clock*, a key clock transcription factor, in individual clock neurons in whole-mount *Drosophila* adult brains (**Figure 1C**). We developed a simple, robust smFISH protocol that takes 3 days from brain dissections to slide preparation (**Figure 1D, 1E**). Next, we generated an accurate three-dimensional segmentation model of clock neurons using machine learning-based approaches by adopting a pretrained decoder-encoder convolutional neural network model. We combined our segmentation analysis and a spot-fitting algorithm, AIRLOCALIZE^24^, into an integrated workflow to acquire cell-specific smFISH data in an automated and reproducible manner (**Figure 1F, Figure 2**). Using these tools, we visualized and quantified *Clock* mRNA distribution in Large ventral lateral clock neurons (lLNvs), one of the main *Drosophila* clock neuron groups.

**Figure 2.**
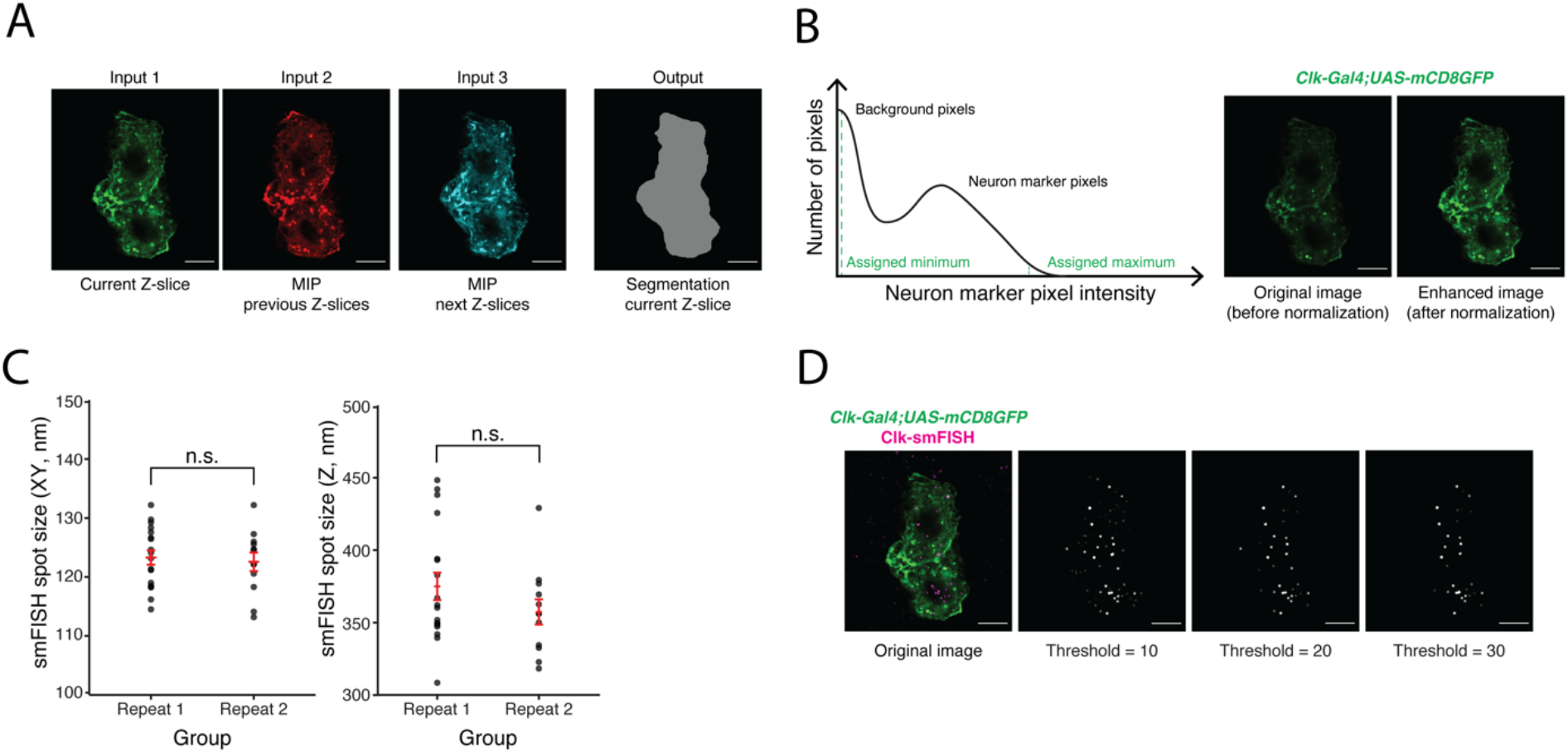
Development of the machine-learning based data analysis pipeline. **A)** Representative images of current and projections of neighboring Z-slices of lLNvs. Maximum intensity projections (MIP) of neighboring Z-slices are used to incorporate three-dimensional information of lLNvs into the segmentation model. **B)** Panel on the left is an illustration of a histogram stretching algorithm. Panel on the right shows representative images of lLNvs before and after normalization. The goal of normalization is to obtain comparable image contrast regardless of original experimental conditions. **C)** Experimentally measured smFISH spot size in XY and Z directions. The values are found to be highly consistent across different biological replicates. **D)** Representative images of detected *Clock* smFISH spots in lLNvs with different threshold settings. Proper threshold selection is important for correct identification of the smFISH spots. Scale bars, 5μm. Statistical test used is unpaired, two-tailed Student’s *t-*tests assuming unequal variances. Individual data points, mean, and s.e.m. (standard error of mean) are shown.

Our results show that *Clock* mRNA levels cycle in lLNvs, consistent with past biochemical studies and single-cell RNA-seq studies^16^, with peak levels at ZT4 (ZT-Zeitgeber time, ZT0-time of lights on, ZT12-time of lights off) with ~80 molecules per neuron and trough levels at ZT16 with ~30 molecules per neuron (**Figure 3, Figure 4**). Consistent with past studies^25^, we show that *Clock* mRNA cycling is abolished in *per*^*01*^ null mutants. Further, we show that clock-neuron specific knockdown of PDP1ε led to consistently low levels of *Clock* mRNA levels throughout the circadian cycle, and knockdown of VRI led to consistently high levels of *Clock* mRNA levels (**Figure 5**). Together, these studies describe a powerful technique to directly visualize and quantify the number and subcellular location of core clock mRNAs in individual neurons, which can open new investigations into the role of spatiotemporal localization of mRNAs in controlling circadian rhythms. The methods described here can be easily adapted to directly visualize gene expression at the single-cell level in *Drosophila* brains.

**Figure 3.**
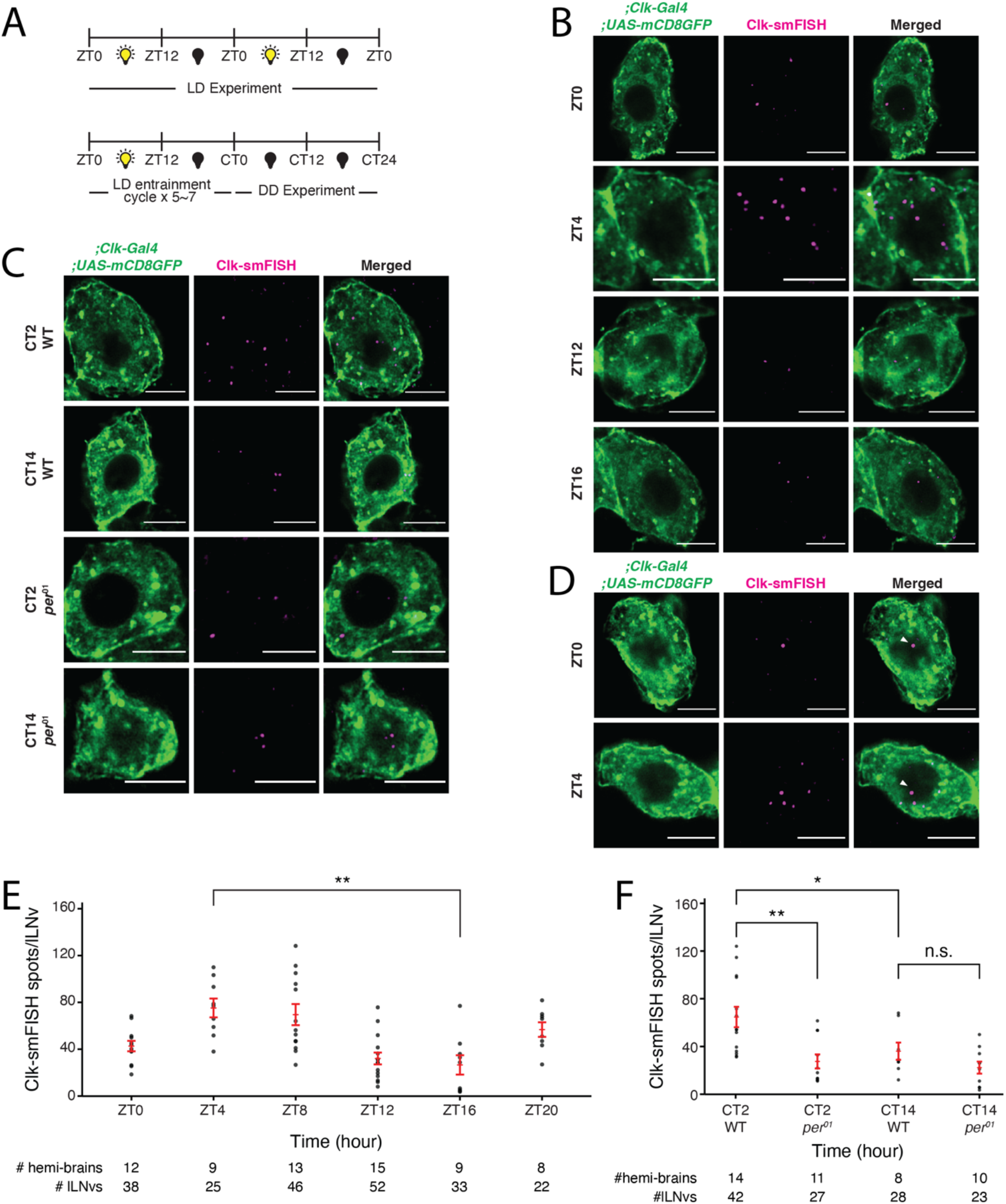
Visualization and quantification of *Clock* mRNA molecules in lLNvs over the circadian cycle. **A)** Entrainment schedule for light-dark (LD) or constant darkness (DD) experiments. We performed *Clock* smFISH experiments using *Clk-GAL4>UAS-CD8GFP* flies under 12h Light: 12h Dark conditions. **B)** Representative images of *Clock* smFISH spots in lLNVs at various timepoints over the circadian cycle. ‘ZT’ refers to Zeitgeber Time; ZT0 denotes the time of lights on and ZT12 denotes the time of lights off. *Clock* mRNA spots are shown in magenta. **C)** Representative images of *Clock* smFISH spots in lLNvs from control and *per*^*01*^ mutant flies in constant darkness conditions. ‘CT’ refers to Circadian Time. *Clock* smFISH spot count is lower at both the timepoints over the circadian cycle in *per*^*01*^ mutants. **D)** Representative image of a single large *Clock* smFISH spot of higher intensity in the nuclei of lLNvs, which corresponds to the active transcription site. **E)** and **F)** show quantification of number of *Clock* smFISH spots per individual lLNvs over the circadian cycle in wild-type (E) and *per*^*01*^ mutant flies (F). Scale bars, 5μm. Statistical test used is unpaired, two-tailed Student’s *t-*test assuming unequal variance. \**P* < 0.05, \*\**P* < 0.005, n.s.-not significant. Individual data points, mean, and s.e.m. (standard error of mean) are shown.

**Figure 4.**
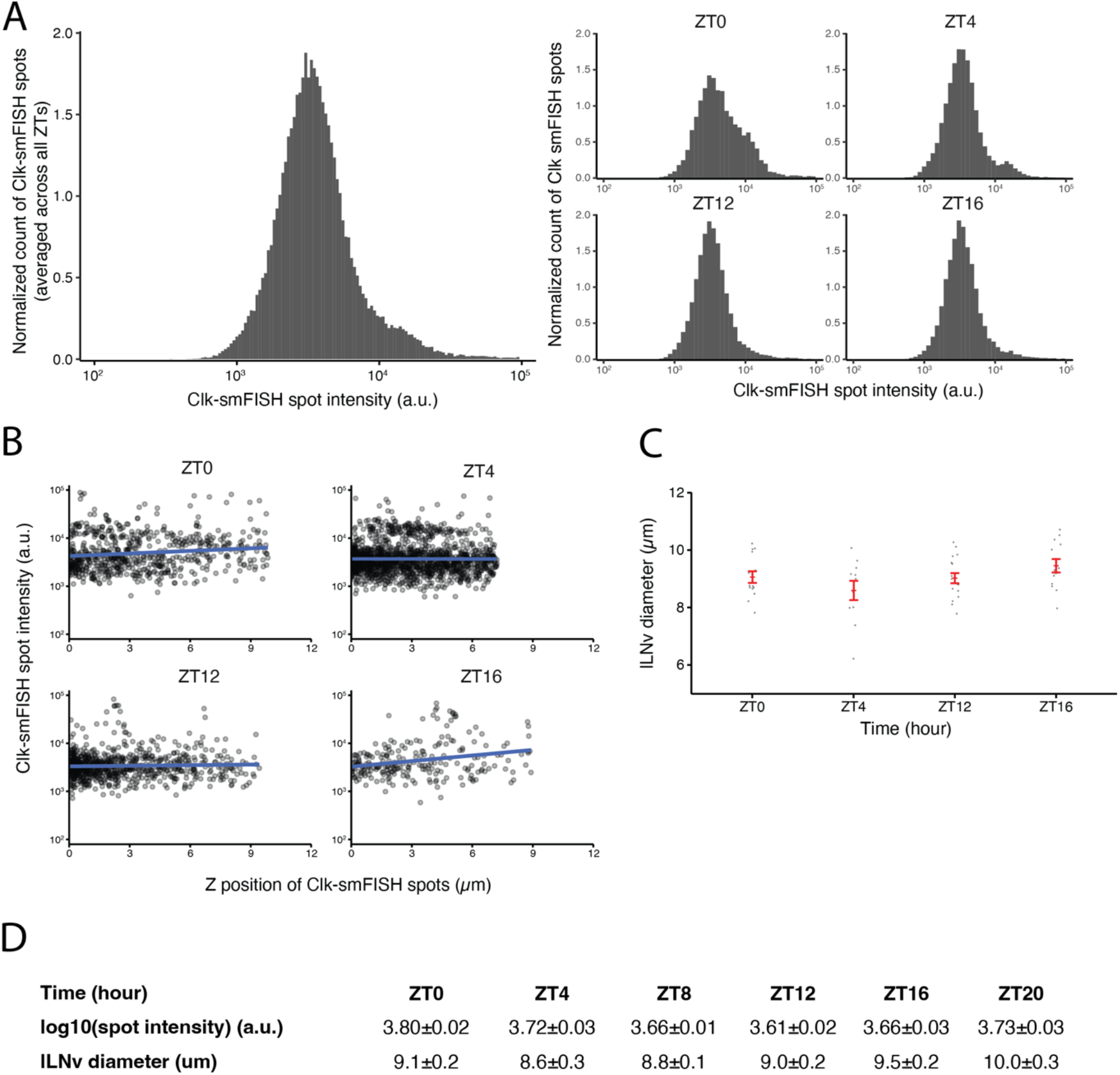
Quality control plots for *Clock* smFISH spots in lLNvs. **A**) Histogram plots of *Clock* smFISH spot intensities in lLNvs. Panel on the left shows averaged data across all ZT’s over the circadian cycle, and panels on the right show data from individual ZT timepoints. **B**) *Clock* smFISH spot intensity distribution across all Z-slices of representative lLNvs at different ZT’s over the circadian cycle. **C)** Equivalent diameters of lLNvs segmentation masks at different ZT’s. **D)** Summary table of the quality control metrics corresponding to *Clock* smFISH spot intensities and lLNv diameters at different ZT’s over the circadian cycle. Individual data points, mean, and s.e.m. (standard error of mean) are shown.

**Figure 5.**
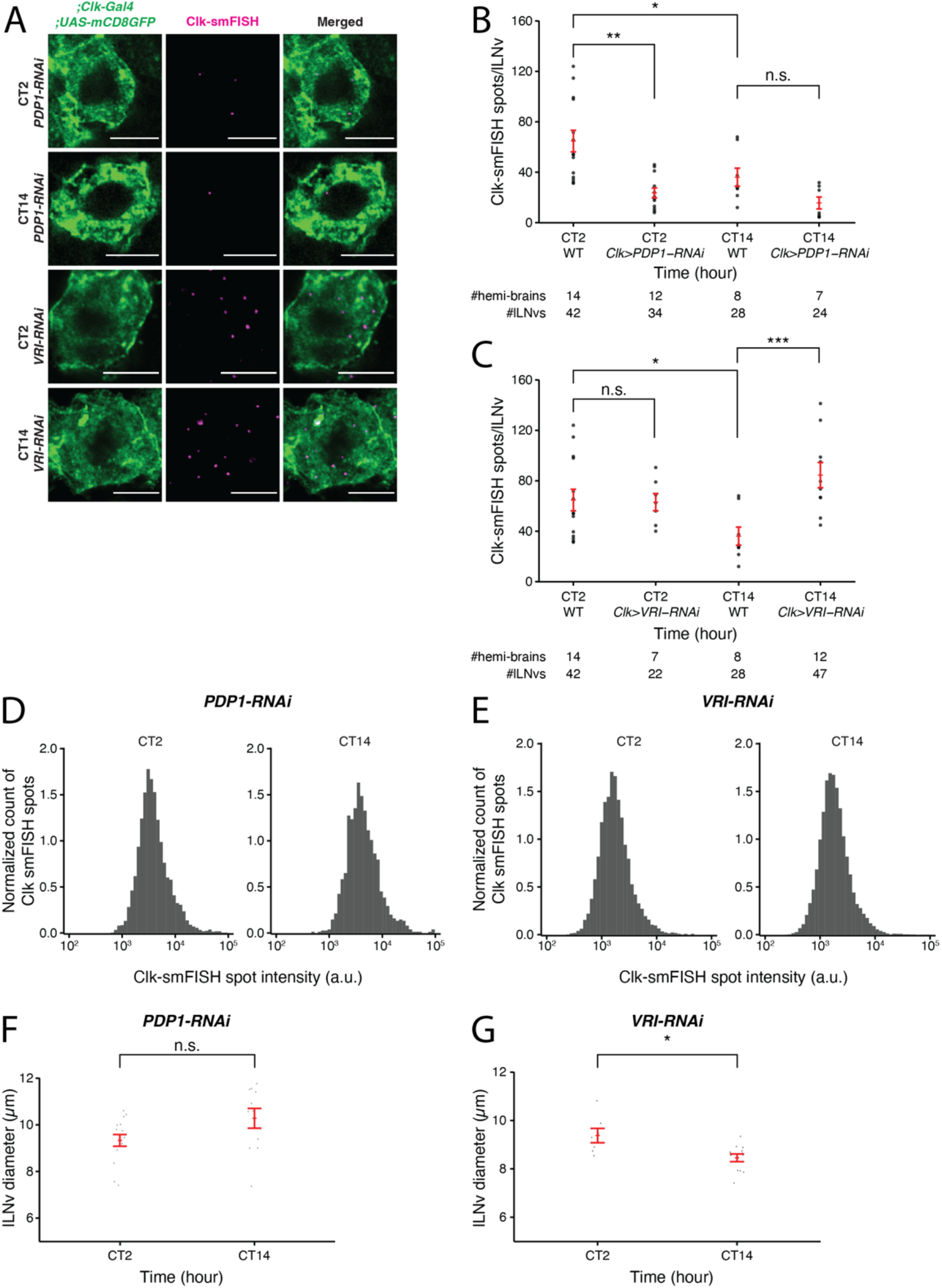
*Clock* mRNA rhythms are disrupted upon clock neuron specific-knockdown of PDP1 and VRI. **A)** Representative images of *Clock* smFISH spots in lLNvs from *Clk-Gal4;UAS-PDP1-RNAi* and *Clk-Gal4;UAS-VRI-RNAi* flies at different CT’s over the circadian cycle. **B, C)** Quantification of the number of *Clock* smFISH spots per lLNv in PDP1-RNAi (B), and VRI-RNAi (C) flies at different CT’s. **D, E)** Histogram plots of *Clock* smFISH spot intensities in lLNvs at different CT’s over the circadian cycle in PDP-RNAi (D) and VRI-RNAi (E) conditions. **F, G**) Equivalent diameters of lLNvs segmentation masks at different CT’s in PDP-RNAi (F) and VRI-RNAi (G) flies. Scale bars, 5μm. Statistical test used is unpaired, two-tailed Student’s *t-*test assuming unequal variance. \**P* < 0.05, \*\**P* < 0.005, \*\*\**P* < 0.0005, n.s.-not significant. Individual data points, mean, and s.e.m. (standard error of mean) are shown.

## Results

To directly visualize the subcellular localization and quantify the number of core clock mRNAs in individual clock neurons, we adapted single molecule RNA-Fluorescence *In Situ* Hybridization (smFISH) method for use in *Drosophila* clock neurons and developed a machine-learning based analysis pipeline to analyze the data. Below, we describe the smFISH method and analysis pipeline in detail and present our findings.

### Procedure overview

Our smFISH procedure consists of five parts: smFISH sample preparation, image acquisition, data analysis, quality control check, and final plot generation (**Figure 1D**). For sample preparation, *Drosophila* brains are dissected, fixed, and hybridized with smFISH probes with the whole protocol taking 3 days requiring little hands-on time (∼1 hour/day) (**Figure 1E**). For imaging, we used a glycerol-based mountant for tissue clearing and acquired images using a high-resolution Airyscan confocal microscope, which enabled us to acquire well defined diffraction-limited spots corresponding to individual mRNA molecules. To automate the process of identification of the cytoplasm and nucleus of lLNvs, a group of clock neurons, we developed a custom convolutional neural network (CNN) segmentation model to generate three-dimensional lLNv clock neuron masks. Performance of our CNN segmentation model was validated by manual inspection of CNN-generated masks on a testing dataset. To quantify the intensity and visualize the subcellular location of individual mRNA spots, we used a published spot detection algorithm, AIRLOCALIZE^24^. Finally, we developed an integrated pipeline in which all data analysis workflows are automated by the workflow manager Snakemake^26^ (**Figure 1F**). After the data analysis step, we generated quality control plots to verify the consistency of our results, identify noisy image outliers, and optimize analysis parameters. The analysis can be rerun with optimized parameters in a fully automated and reproducible manner. In the next subsections, we describe individual steps of the protocol in detail.

### Fluorescent oligonucleotide probes for smFISH experiments

To enable detection of *Clock* mRNA transcripts, we ordered commercial fluorescent oligonucleotide probes from Biosearch Technologies, Inc. First, we designed ∼48 20nt short oligonucleotide probes that span across the full-length of all annotated *Clock* mRNA isoform species covering all exons using the online Stellaris RNA-FISH Probe Designer (**Figure 1C**). For genes with alternative splicing isoforms, shared exons can be used to design smFISH probes to ensure detection of all isoforms. The designer provides masking of repetitive sequences to improve probe specificity, and we generally start with the most stringent masking level. In our experience, thirty probes are generally adequate, but the minimum number of probes that can yield high signal-to-noise ratio images is likely dependent on microscope setup (*e*.*g*., objective numerical aperture, detector sensitivity etc). If the target mRNA is short or contains repetitive sequences, it might not be possible to design ∼48 smFISH probes targeting the mRNA. In such a case, masking level can be adjusted to increase the number of smFISH probes. Each of the 20nt probe is conjugated at the 5’ end to a fluorophore and collaborative binding of ∼48 probes to an individual mRNA molecule enables detection of a single mRNA spot by fluorescence microscopy. Upon receipt of the probe pool, we prepare aliquots of 20mM probe stock in 10mM Tris-Cl, pH 8.5 and store them in −20C.

### Sample preparation for smFISH experiments

In our smFISH experiments, we typically use transgenic flies in which cell membranes are fluorescently labeled. Specifically, we drive the expression of CD4-tdTomato or CD8-GFP^27^ exclusively in clock neurons by crossing them with pan-clock neuron driver flies, *Clk856-GAL4*^28^. On day 1, *Drosophila* brains are dissected, fixed in 4% paraformaldehyde, and permeabilized in 70% ethanol at 4ºC overnight. Fixation is a crucial step in smFISH experiments, as both under-fixation as well as over-fixation of tissues can result in samples in which cellular morphology is not well preserved. Permeabilization with 70% ethanol is another necessary step to allow the smFISH probes to pass through the cell membrane and enter the cell. On day 2, brains are washed with PBS and then incubated in hybridization solution with probes overnight at 37ºC. On day 3, brains are washed with PBS to remove the hybridization solution and mounted on a slide using a glycerol-based mountant for tissue clearing (see Methods section for a detailed protocol). This smFISH protocol is robust, simple to implement, taking 3 days from brain dissections to slide preparation, and produces highly reproducible results across experimental repeats (**Figure 1E**).

### Microscope setup for image acquisition

The goal is to obtain clock neuron images where smFISH spots and the neuron cell membrane marker are clearly visible and can be distinguished from the background. In general, image acquisition parameters should be optimized taking into account both the clock neuron marker and smFISH probe channels, and the same settings need to be reused across all experimental conditions and repeats to ensure data consistency. Filter settings must be compatible with the fluorophore used and appropriate laser power and detector gain should be determined. While larger laser power and detector gain will yield higher pixel intensities, high laser power will lead to significant bleaching during acquisition and high detector gain will lead to increased random noise. Finally, to further suppress random noise, pixel averaging can be performed to obtain distinct diffraction-limited spots in the smFISH probe channels and clear cell boundaries in the neuron marker channel across all Z-slices.

### Segmentation of clock neurons

The goal of the segmentation model is to assign a classification label to each pixel of the Z-stack image, i.e., to determine whether a pixel can be assigned to a ‘clock neuron’ or to the background. The clock neuron marker used in our studies, CD8-GFP, not only labels the plasma membrane but also the components of the secretory pathway intracellularly. Therefore, CD8-GFP cell marker intensity varies significantly throughout the cytoplasm, making conventional segmentation methods such as simple thresholding (*e*.*g*., Otsu method^29^) ineffective. To solve this issue, we adopted a publicly available pretrained encoder-decoder convolutional neural network (CNN) model based on the U-Net architecture^30^ for semantic segmentation of clock neurons. In short, the CNN model computes complex features based on the original clock neuron marker signal (*i*.*e*., encoding) and then assigns classification labels based on the computed features (*i*.*e*., decoding). Publicly available pretrained CNN models are typically developed with two-dimensional image datasets^30^. To provide an accurate segmentation model of 3-dimensional clock neurons, for each Z-slice we input the following images to the CNN model: 1) current Z-slice, 2) a maximum intensity projection (MIP) of neighboring Z-slices within 0.4μm in one direction, 3) a MIP of neighboring 0.4μm slices in another Z-direction. (**Figure 2A**).

Further, during model training, we found that CD8-GFP intensity level may vary significantly across different experimental conditions, which hindered the performance of the model. To address this issue, we adopted a MATLAB-based histogram stretching algorithm^31^ to normalize the levels of CD8-GFP intensities among different images. Briefly, the distribution of cell marker pixel intensities follows a bimodal distribution; most pixels will show near-zero values (background pixels) while some show significantly higher values (true CD8GFP signal) (**Figure 2B, left panel**). We then take the 2% dimmest and brightest intensities and assign them to be the minimum and maximum values. Figure 2B panel on the right shows the original image, in which CD8-GFP signal is dim, and the enhanced image after normalization.

One of the caveats to adopting complex CNN models to generate segmentation masks is the limited training set size which may lead to suboptimal models. To efficiently train our CNN image segmentation model, we augmented (i.e., increased the size of) our training dataset by computationally generating images with varying brightness levels from the original training data set^32^. We note that lLNv clock neurons are typically contiguous to each other, which makes it difficult to identify individual lLNvs using our segmentation model. In fact, our segmentation model provides an outline of all the lLNvs in each hemi-brain (**Supplementary Video 1**). We verified the performance of the segmentation model by visual inspection of testing datasets, and we were able to generate segmentation masks of the cytoplasm and the nucleus of lLNvs with ~10 manually annotated Z-stacks (**Supplementary Video 1**).

### smFISH spot detection

We adopted the publicly available spot counting package, AIRLOCALIZE^24^, for mRNA spot detection and quantification. In short, diffraction-limited spots are modeled as a three-dimensional gaussian function with lateral (i.e., XY) symmetry. Parameters of the gaussian function, essentially sizes of the spot in both XY and Z directions, can be measured using a graphical interface provided by the AIRLOCALIZE package. Using our Airyscan confocal microscope, we found that the spot sizes in both XY (∼120 nm) and Z (∼400 nm) directions are highly reproducible (**Figure 2C**).

To detect smFISH spots in an image, the AIRLOCALIZE algorithm first applies a global threshold to reject noise in the image. Then, the algorithm finds candidates of smFISH spots by identifying locally brightest pixels. Finally, gaussian fitting is performed using the identified candidate spot locations and a list of ‘true’ smFISH spots is obtained. In Figure 2D, we show the effect of applying different threshold settings on smFISH spot detection. In this example image, the number of detected spots in lLNvs changes with threshold setting. At the minimum threshold tested, we were able to detect ∼70 smFISH spots. At the maximum threshold tested, ∼30 spots were detected (**Figure 2D**). The global threshold therefore affects smFISH spot detection results and should be set empirically. Typically, we find that the same threshold can be used for all images acquired in one experimental batch. Moreover, our analysis pipeline (discussed in the next section) provides spot visualization capabilities, which can be used to optimize the threshold parameter (**Supplementary Video 2**).

### Data preprocessing and integration of the workflows

To ensure reproducibility and aid optimization of analysis parameters, we developed an automated analysis pipeline using the Snakemake workflow management system^26^ (**Figure 1F**). First, the raw data is converted to a lossless, open-source OME-TIFF format by the Bio-Formats package^33^. Next, smFISH and cell marker images are used for spot detection and cell segmentation, respectively. To obtain the list of smFISH spots within clock neurons, we only consider smFISH spots whose centroids overlap with the lLNv segmentation masks. Finally, quality control plots and images of all intermediate outputs (segmentation masks and detected smFISH spots within clock neurons) are generated together with the final output plots. In the final output plots, we report the number of detected smFISH spots per clock neuron and the locations and intensities of smFISH spots (**Figure 1F**). Quality control plots are discussed in more detail in the following section. The data analysis pipeline with detailed documentation is publicly available online at Github repository (https://github.com/yeyuan98/rna_fish_analysis).

### Quality control check

As a large number of Z-stack images are generated in each experiment, unbiased consistency verification and identification of noisy images are vital for our workflow. Towards this end, we developed a series of quality control plots (for an overview see **Figure 1F**, for representative results see **Figure 4**). First, we generated intensity histogram plots of all detected smFISH spots in individual images (**Figure 4A**). As each mRNA molecule is bound by a similar number of fluorophores, intensities of individual smFISH spots should be consistent among different images and experimental conditions. Second, to determine whether working distance is adequate, smFISH spot intensity distribution across all Z-slices is plotted for each image with a linear fit (**Figure 4B**). If the objective working distance is sufficient, an approximately constant spot intensity distribution across all Z-slices is to be expected. Finally, to determine the performance of our segmentation model, physical volumes of the segmentation masks are computed and summarized as equivalent diameter values, which is expected to be ∼10 microns for all lLNVs (**Figure 4C**).

### Visualization and quantification of *Clock* mRNAs in lLNv neurons

Using our smFISH method and data analysis pipeline, we performed smFISH experiments to detect *Clock* mRNA transcripts in large ventral lateral neurons (lLNvs) in whole-mount *Drosophila* brains across the circadian cycle. We entrained the flies to 12h Light: 12h Dark cycles (‘ZT0’ refers to time of lights on and ‘ZT12’ refers to time of lights off) and performed smFISH experiments at various timepoints across the circadian cycle (**Figure 3A**). We observed ∼80 *Clock* mRNAs per lLNv at ZT4 (peak) and ∼30 mRNAs per lLNv at ZT16 (trough) (**Figure 3B, Figure 3E, Supplementary Figure S1**), consistent with past qPCR and single-cell RNA-seq studies^16^. We also found that *Clock* mRNA transcript numbers do not cycle in *per*^*01*^ null mutant flies^34^ and remain low throughout the circadian cycle (**Figure 3C, Figure 3F, Supplementary Figure S2**). These results are consistent with past studies^25^ which reported that CLK protein fluorescence levels are lower in *per*^*01*^ mutant flies compared to control flies. Further, we tested how *Clock* mRNA levels are affected upon clock neuron-specific RNAi-knockdown of PDP1 and VRILLE, transcriptional regulators of Clock. Our smFISH results demonstrate that knockdown of PDP1 abolished *Clock* mRNA rhythms with lower number of *Clock* mRNA molecules at different timepoints over the circadian cycle, consistent with PDP1’s function as a *Clock* activator (**Figure 5A, Figure 5B**). As expected, knockdown of VRILLE, which is a transcriptional repressor of *Clock*, led to a higher number of *Clock* mRNA molecules over the circadian cycle (**Figure 5A, Figure 5C**).

Quality control is an essential component of our data analysis pipeline, enabling us to verify the validity of our experimental results. As individual mRNAs are expected to be bound by a similar number of fluorescent oligonucleotide probes, the histogram plots of smFISH spot intensities are expected to show predominantly a single peak with consistent distribution among different samples. The individual and merged intensity histogram plots generated from our experimental data show that *Clock* smFISH signal intensities in lLNvs from wild-type flies are highly consistent (**Figure 4A**). Next, as we usually scan ∼10μm in the Z-axis for imaging lLNvs in a hemi-brain, we examined how smFISH spot intensity varies across each spot’s Z position in each acquired image (**Figure 4B, Figure 4D**). Again, as individual mRNAs are bound by a similar number of fluorescent oligonucleotide probes, if working distance is sufficient smFISH spot intensity should not depend on Z-position. Indeed, >80% of our acquired images show no significant correlation between smFISH spot intensity values and Z position of the spots. Finally, equivalent diameters of segmentation masks of lLNvs demonstrate consistent volumes expected for lLNv neurons (**Figure 4C, Figure 4D**). With these quality controls, we were able to confirm the consistency and validity of our experimental data. Intermediate results of the pipeline including segmentation masks and identified smFISH spots are readily available for visual inspection (**Supplementary Videos 1, 2**).

## Discussion

Here, we present a streamlined single molecule RNA-fluorescent in situ hybridization (smFISH) protocol and a machine-learning based data analysis pipeline that enables direct visualization of the subcellular location of individual mRNAs and precise quantification of the total number of mRNA molecules in clock neurons. Using this method, we show that *Clock* mRNAs cycle in lLNvs with peak levels (~80 molecules/lLNv) at ZT4 and trough levels (~30 molecules/lLNv) at ZT16. We validated our smFISH experimental protocol and data analysis pipeline by analyzing *Clock* mRNA levels in wildtype and mutant conditions.

In addition to transcriptional regulation, post-transcriptional regulation plays a prominent role in regulating gene expression. Localization of mRNAs to specific subcellular structures (*e*.*g*., P-bodies, stress granules, nuclear speckles) has been shown to be vital for post-transcriptional regulation in many different biological processes^35-37^. To visualize mRNA localization, RNA fluorescence *in situ* hybridization (RNA-FISH) methods have been widely adapted in the past in many model systems^38-40^. In general, RNA-FISH technique uses a number of oligonucleotide probes that can specifically bind to specific sequences of a mRNA, and direct detection (as shown in our *Clock* smFISH experiments) or amplification-based methods (*e*.*g*., hybridization chain reaction, HCR^41^) can be used to visualize the mRNAs. While amplification-based methods offer more flexibility with regard to target length and yield better signal-to-noise ratios, direct detection methods generate individual diffraction-limited spots for each single mRNA molecule with well-defined intensity and shape profiles.

smFISH protocol requires hybridization of tens of ∼20nt oligonucleotides to the same RNA molecule of interest. Therefore, a relatively long transcript (∼1.5 kb) is required for adequate signal-to-noise ratio (SNR). If short transcripts such as individual exons or introns need to visualized, amplification-based methods such as hybridization chain reaction (HCR) may be adopted^41^. However, while spatial information is typically retained in amplification-based methods, precise quantification of the number of mRNA molecules might require optimization as signal amplification kinetics must be determined empirically. While smFISH and HCR protocols involve fixing the sample and hybridizing it with fluorescent oligonucleotide probes, MS2-MCP system enables live imaging of individual mRNA molecules with high temporal and spatial resolution^24,42^. Briefly, RNA loops derived from the bacteriophage MS2 can be inserted in the 3’UTR of an mRNA of interest and co-expressed with MCP, which is a MS2 coat protein that can tightly bind to MS2 binding sites, fused to fluorescent proteins. While the MS2-MCP system enables live imaging of individual mRNA molecules, however, addition of stem-loops to the mRNA molecule of interest may affect its function and/or transcription/degradation kinetics. In conclusion, our smFISH method and the data analysis pipeline presented here is a powerful tool that can be employed to study subcellular organization of any clock mRNA molecule in *Drosophila* clock neurons.

## Materials and Methods Data reporting

No statistical calculations were used to pre-determine sample sizes. Samples were not randomized for the smFISH experiments and blinding was not used during the data analysis step. We reported the sample sizes for each of our experiments in the figures.

### *Drosophila* strains

Flies were raised on cornmeal-yeast-sucrose food under a 12:12 Light:Dark cycle at 25°C and 60-70% humidity. The following flies used in the study were previously described or obtained from the Bloomington Stock Center: *Clk-GAL4*^28^ (expressed in all clock neuron classes), *per*^*01* 34^, *UAS-CD8GFP*^27^, *UAS-Pdp1-RNAi* (BL-40863), *UAS-vri-RNAi* (BL-40862). Specific genotypes of flies used in the experiments are presented in the figures.

### Solutions used for smFISH

Prepare all solutions with RNase-free tubes and reagents and filtered pipette tips. Aliquot reagent stocks to reduce chance of contamination.

*Fixation solution*: 4% formaldehyde in 1X phosphate-buffered saline (PBS) and is made from 16% EM-grade formaldehyde (Electron Microscopy Sciences 15710) and 10X PBS (Invitrogen AM9625) in RNase-free water.

*Permeabilization solution*: 70% ethanol diluted from 200 proof ethanol (Thermo Scientific T038181000) in RNase-free water.

*Hybridization solution*: 2X saline-sodium citrate (SSC, Invitrogen AM9763), 10% dextran sulfate (Thermo Scientific AC433240050), 1mg/mL E.coli tRNA (Roche 10109541001), 2mM ribonucleoside vanadyl complex (RVC, New England Biolabs S1402S), 0.5% bovine serum albumin (BSA, Roche 10711454001), 10% formamide (Sigma, F9037), and 5μM oligonucleotide probes conjugated to fluorophores (Stellaris, LGC Biosearch Technologies) in RNase-free water.

*Wash solution*: 2X SSC and 10% formamide in RNase-free water.

We prepare 20% (weight/volume) dextran sulfate stock from powder and store at 4ºC. Fixation and permeabilization solutions are prepared fresh on day of experiment. RVC and formamide aliquots are used one-time once thawed.

### smFISH protocol

1. Prepare fixative solution. For each sample, put 500uL of fixative to a well of a clean glass dish and leave on ice.
2. Dissect at least 10 brains in 1X PBS for each sample. Put dissected brains directly into the fixative on ice in the glass dish. Dissections should be completed within 30 minutes.
3. Place the glass dish on a nutator and incubate for 20 minutes at room temperature.
4. Remove the fixative and perform three quick rinses with 1X PBS.
5. Remove solution and wash twice with 1X PBS, 5 minutes per wash at room temperature with nutation.
6. Remove 1X PBS completely and add 500uL of permeabilization solution to the glass dish. With a P1000 tip, carefully transfer the brains into a protein low-bind tube. Avoid damaging the brains during transfer.
7. Fill the tubes with extra permeabilization solution to a final volume of 1mL. Incubate the sample overnight at 4ºC with nutation, protected from light.
8. Prepare hybridization solution, 100μL per sample and leave on ice protected from light. Prepare wash solution, 1mL per sample and leave at room temperature. Hybridization solution is viscous and should be well-mixed by pipetting (>10 times).
9. Aspirate permeabilization solution and add 1mL of wash solution. Incubate 5 minutes at room temperature with nutation.
10. Aspirate the wash solution completely and add 100μL of hybridization solution containing probe(s). Gently pipette to make sure all brains are collected in the solution.
11. Incubate overnight in a 37ºC incubator, protected from light.
12. Add 1mL of wash solution without removing the hybridization solution. Gently pipette once to mix without touching the brains. Incubate at 37ºC for 30 minutes.
13. Aspirate off the solution and wash one more time with 1mL of wash solution at 37ºC for 30 minutes. Supplement the wash solution with 1μg/mL DAPI for nucleus staining.
14. Remove the wash solution and mount the samples in Prolong glass mountant. Make sure the brains are mounted in the appropriate orientation to visualize the clock neurons. Seal the samples immediately after mounting and leave at room temperature overnight protected from light.
15. The samples are ready for imaging the next day.

### Imaging protocol

Flies were entrained to Light-Dark (LD) cycles with lights on for 12 hours and off for 12 hours for 5-7 days, and then released into complete darkness (DD) for 6-7 more days. Zeitgeber Time (ZT) 0 marks the time of lights on, and ZT12 marks the time of lights off. Circadian time (CT) refers to times in complete darkness (DD) and CT0 is the start of the subjective light phase and CT12 is the start of the subjective dark phase, the times when the light transitions would have occurred had the LD cycle continued.

All flies used for smFISH experiments were placed in density-controlled food vials (4 females and 4 males) and entrained for 5-7 days in incubators. We performed all our smFISH experiments on 5-7 day old male or female flies, and did not notice any differences in our experimental results. The fly genotype and ZT was as described in the figure legends. We used the GAL4/UAS system to express transgenes in clock neurons in the brain. We imaged our samples using a Zeiss LSM800 laser scanning confocal microscope with AiryScan super-resolution module (125 nm lateral and 350 nm axial resolution). We have acquired our images using a 63x Plan-Apochromat Oil (N.A. 1.4) objective and 405, 488, and 561, and 647 nm laser lines.

### Statistical tests

All experiments were repeated at least twice independently. All statistical significance tests presented are unpaired, two-tailed Student’s *t-*tests assuming unequal variances.

## Data Availability Statement

The analysis pipeline is publicly available online at https://github.com/yeyuan98/rna_fish_analysis.

## Author Contributions

Ye Yuan, Conceptualization, Data curation, Formal analysis, Validation, Investigation, Visualization, Methodology, Writing—original draft, Writing—review and editing; Marc-Antonio Padilla and Dunham Clark, Formal analysis, Visualization; Swathi Yadlapalli, Conceptualization, Investigation, Supervision, Funding acquisition, Methodology, Writing— original draft, Writing—review and editing.

## Acknowledgments

The authors acknowledge help from Dr. Yukiko Yamashita and Dr. Jaclyn Fingerhut in adapting smFISH protocol for use in adult *Drosophila* brains. The authors also thank Yangbo Xiao for aiding in the preparation of the figures. This work was supported by funds from the NIH (grant no. R35GM133737 to S.Y.), Alfred P. Sloan Fellowship (to S. Y.) and McKnight Scholar Award (to S. Y.).

---

**Supplementary Figure 1.**
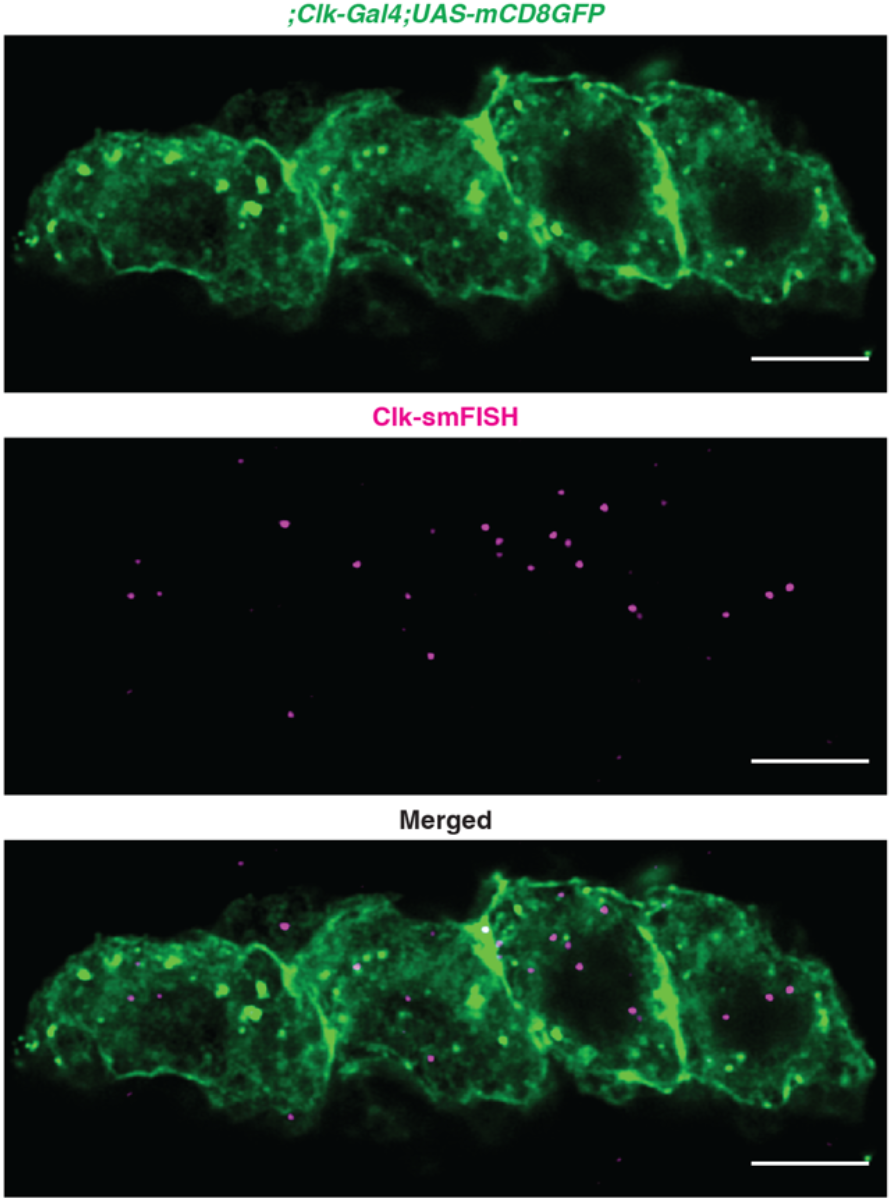
Zoomed-out representative image of *Clock* smFISH spots in all lLNvs. Here we show a representative Z-slice of all four lLNvs observed in a hemi-brain at ZT4. Scale bars, 5 μm.

**Supplementary Figure 2.**
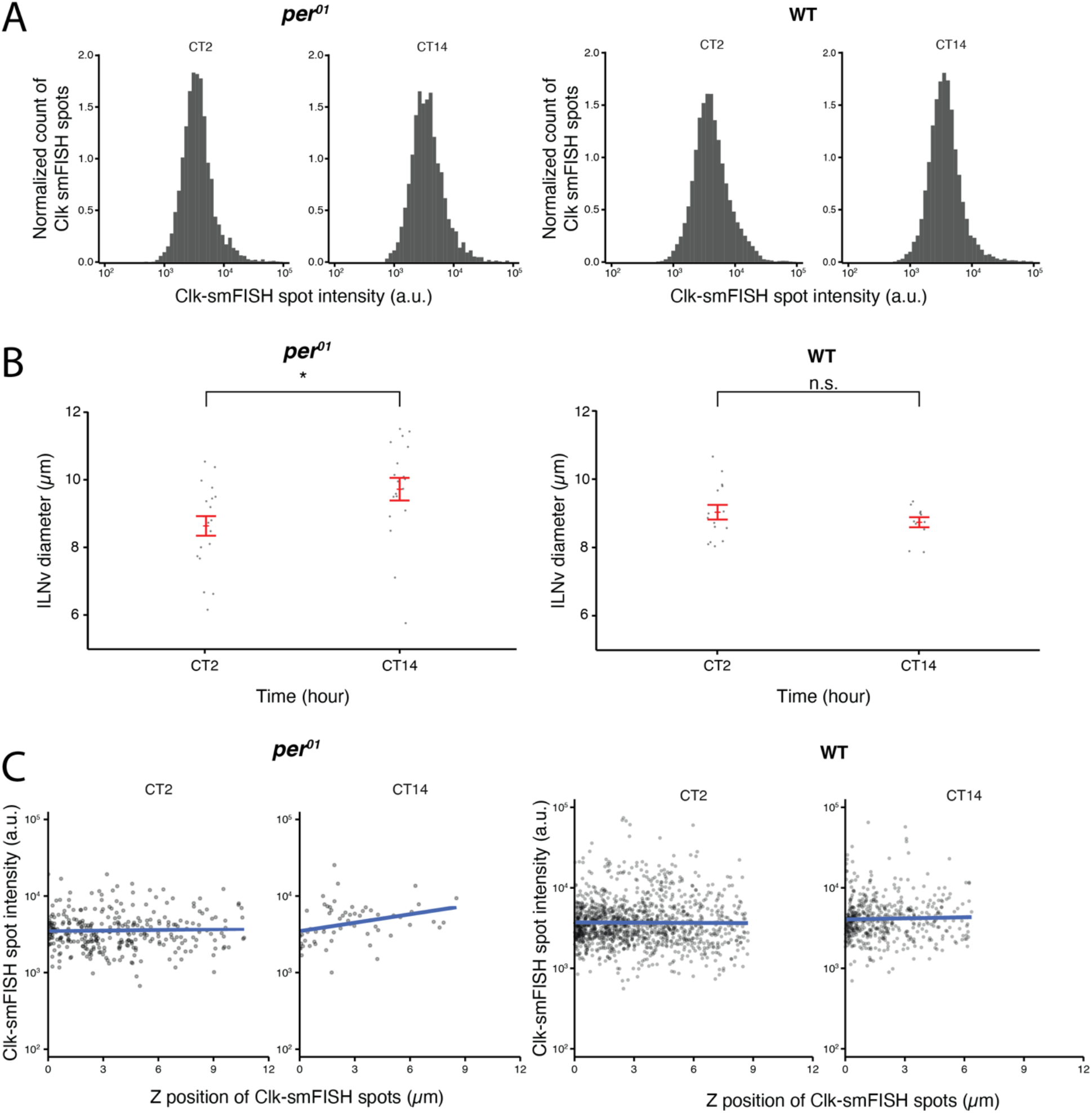
Quality control plots for *Clock* smFISH spots in lLNvs from *per*^*01*^ null mutant flies. **A**) Histogram plots of *Clock* smFISH spot intensities in lLNvs from *per*^*01*^ (left) and wildtype (right) flies at different CT’s over the circadian cycle. **B**) Equivalent diameters of lLNvs segmentation masks from *per*^*01*^ (left) and wildtype (right) flies at different CT’s. **C**) *Clock* smFISH spot intensity distribution across all Z-slices of representative lLNvs from *per*^*01*^ (left) and wildtype (right) flies at different CT’s. Statistical test used is unpaired, two-tailed Student’s *t-*test assuming unequal variance. \**P* < 0.05, n.s.-not significant. Individual data points, mean, and s.e.m. (standard error of mean) are shown.

**Supplementary Video 1. Representative segmentation mask of lLNv neurons**. A Z-stack video showing a group of 4 lLNv clock neurons marked with CD8-GFP (left panel) and representative segmentation masks from our CNN model (right panel). Scale bar: 4 μm.

**Supplementary Video 2. Visualization of detected *Clock* smFISH spots in lLNvs**. A Z-stack video showing *Clock* mRNA probe signals in magenta (Quasar 670 fluorophore) and clock neurons in green (CD8-GFP) in the left panel. Right panel shows visualization of detected *Clock* smFISH spots in lLNvs using AIRLOCALIZE algorithm. Scale bar: 4 μm.

